# Profiling of *Zea mays* L. proteome at early stages of compatible interactions with *Meloidogyne arenaria* indicates changes in signaling, oxidative stress responses, and S-adenosylmethionine biosynthesis

**DOI:** 10.1101/2023.06.28.546826

**Authors:** Przybylska Arnika, Wrzesińska-Krupa Barbara, Obrępalska-Stęplowska Aleksandra

**Affiliations:** Department of Molecular Biology and Biotechnology, Institute of Plant Protection – National Research Institute, Poznań, Poland

**Keywords:** *Meloidogyne arenaria*, *Zea mays*, root-knot nematodes, defense response, cell wall modifications, signal transduction, oxidative stress, S-adenosylmethionine

## Abstract

Root-knot nematodes (RKNs) are distributed worldwide group of plant parasitic nematodes, with a very wide host range, including mono- and dicotyledonous hosts. *Meloidogyne arenaria* is, next to *M. hapla, M. incognita*, and *M. javanica*, one of the most economically important species from this genus. RKNs during parasitism hijack host metabolism to establish giant cells and to break down plant defense mechanisms. To date, studies on the interaction of RKN with maize (*Zea mays* L.) have been underrepresented, and a description of the early response to attack by these nematodes, vital to understanding the process, is scarce in the literature. We hypothesize that in the early stage of maize response to *M. arenaria* infection, significant changes in the accumulation level of proteins mainly related to plant defense response, plant cell wall modifications, and phytohormone biosynthesis can be observed.

In this study, a mass spectrometry approach and a label-free quantification technique were used to assess the qualitative and quantitative composition of proteins changes in the proteome of maize roots after *M. arenaria* infection. We used a susceptible maize variety and carried out analyses of plant proteome at two time points: 24 hours after nematode inoculation and 3 days after inoculation. Statistical analyses of significant differences between protein intensities were performed for the datasets obtained from healthy and *M. arenaria*-infected plants, and differentially expressed proteins (DEPs), with both lower and higher abundance were determined. DEPs were mapped, classified to the gene ontology (GO) terms into functional categories, and assigned to appropriate Kyoto Encyclopedia of Genes and Genomes (KEGG) processes and pathways.

As a result, a total of 3,743 proteins were identified with 124 DEPs at 24 hpi and 66 at 3 dpi, and significant changes in the accumulation of proteins associated with processes such as cell wall modifications, reaction to stress, as well as processes and pathways related to phenylpropanoid biosynthesis and metabolism, signal transduction and to S-adenosylmethionine biosynthesis.

## Introduction

Root-knot nematodes (RKNs, *Meloidogyne* spp.) are plant parasitic nematodes, with more than 100 described species distributed worldwide (Ye et al. 2019). *Meloidogyne arenaria*, next to *M. hapla, M. incognita*, and *M. javanica*, is among the most damaging species to crops, with a very wide range of hosts, including mono- and dicotyledonous plants (CABI 2022). On the other hand, one of its monocotyledonous hosts, maize, belongs to the world’s major crop species and is an important source of food for humans and animals (Hallauer and Carena 2009). During the plant invasion, RKNs penetrate the root system and hijack the plant metabolism to establish giant cells and root galling (Kyndt et al. 2014).

There are many mechanisms to respond to pathogens’ attacks in plants. Jones and Dangl (2006) described a four-phased “zigzag” model, to explain the plant defense response. According to this model, there are two crucial mechanisms of immunity. The first one is called pathogen-associated molecular patterns (PAMPs)-triggered immunity (PTI), while the second one is an effector-triggered immunity (ETI). The first layer of response is PTI, active via plant pattern-recognition receptors (PRRs) activated by PAMPs, which allows for further plant colonization by RKNs during compatible interactions. When PTI is suppressed by pest effectors, effector-triggered susceptibility (ETS) is induced, but an effector may be recognized by the appropriate proteins in plants activating ETI response. Successful recognition of pathogen effector in ETI leads, in consequence, to the development of hypersensitive response (HR) in infected plant cells and/or systemic acquired resistance (SAR) in the whole plant (Jones and Dangl 2006). PTI response induced in plants by PAMPs goes through signal transduction mechanisms, including the activation of mitogen-activated protein kinases (MAPKs) (McNeece et al. 2019), generation of reactive oxygen species (ROS) (Jwa and Hwang 2017), and activation of the jasmonic acid (JA) and the salicylic acid (SA) signaling pathways (Manosalva et al. 2015). During host response to nematode infection, the JA-related pathway leads to the induction of accumulation of, among others, pathogenesis-related genes (*PR*), such as *PR3* and *PR4* (Hamamouch et al. 2011; Macharia et al. 2020) while the SA-related pathway induces the accumulation of SA signature genes i.e. *PR1, PR2*, and *PR5* (Ali et al. 2018; Hamamouch et al. 2011; Przybylska et al. 2018).

Thus far, many response mechanisms have been described in the compatible interactions between RKNs and their hosts, mostly for dicotyledonous plants but for monocotyledonous are some studies as well (Przybylska and Obrępalska-Stęplowska 2020). The monocotyledonous host for which the response to infection by *Meloidogyne* spp. has been best studied so far is rice. It was found that in the compatible interactions between this plant and *M. graminicola*, mainly JA-related pathway was activated (Nahar et al. 2011). There are also reports that both, JA- and SA-related pathways were induced in the early stage of RKN infection (Kumari et al. 2016; Kumari et al. 2017; Petitot et al. 2017). Additionally, results of proteomic studies, described by Xiang et al. (2020) indicated significant downregulation of proteins related to the alpha-linolenic acid metabolism, phenylpropanoid biosynthesis, glutathione metabolism, and proteins related to plant-pathogen interaction pathways as e.g., calcium/calmodulin-dependent protein kinase in rice variety sensitive to *M. graminicola* infection. The response of maize to *Meloidogyne* spp. attack has been the subject of only a few studies. In our previous research on maize-*M. arenaria* interaction, we observed suppression of defense-related genes associated with both, the JA-pathway and the SA-pathway in the early stage of *M. arenaria* infection (Przybylska et al. 2018). Moreover, Gao et al. (2020) described an increased level of JA in the roots and an increased susceptibility to *M. incognita* in maize mutants lacking lipoxygenase (ZmLOX3). Moreover, in another study, we reported significant upregulation of the gene encoding polygalacturonase in the early stage, while suppression of expression of this gene in later stages of *M. arenaria* infection of susceptible variety of maize (Przybylska and Spychalski 2021). Recently, we also described the first candidate effector protein of *M. arenaria –* MaMsp4, interacting with host proteins involved in plant defense (associated with polyamine and phosphatidylethanolamine biosynthesis) as well as in plant cell wall modification (Przybylska et al. 2023).

Although both, *Meloidogyne* and its host – maize, are economically important as pests and staple food crops worldwide, there are few data showing changes in the proteome of this plant, as well as other monocotyledonous plants, attacked by this group of nematodes. This study aimed at analyzing a proteomic profile of maize during the early stage of compatible interactions with *M. arenaria*. We hypothesize that in the early stage of maize response to *M. arenaria* infection, significant changes in the accumulation of proteins related mostly to plant defense response, plant cell wall modifications, and phytohormone biosynthesis may be observed.

## Material and methods

### Material and samples collection

Proteomic analyses were performed on a variety of maize (PR39F58) with high susceptibility to *M. arenaria* infection, evaluated during our previous study (Przybylska et al. 2018). *M. arenaria* population came from the Institute of Plant Protection – National Research Institute collection. *M. arenaria* larvae were hatched from eggs extracted from maize roots using NaOCl, according to the technique described by Hussey (1973).

Maize plants were grown in a greenhouse at constant day/night temperatures of 25 °C/20 °C and under controlled light conditions. The day/night temperatures were switched at 7 am and 7 pm. The 3–4 week old seedlings at the 4–5 leaves stage were infected with 1500 specimens of *M. arenaria* in J2 invasive stage. Root samples were collected at two time points: 24 hours post inoculation (hpi) and 3 days past inoculation (3 dpi) from three infected plants as well as three healthy plants as a control.

### Proteins extraction and samples preparation

Total proteins from root samples were extracted using the technique described by Frąckowiak et al. (2022). Briefly, the proteins’ pellet was dissolved in 50 μl lysis buffer containing 5% sodium dodecyl sulfate (SDS) and 50 mM triethylammonium bicarbonate (TEAB), pH 8.5. Next, proteins were reduced by the addition of 15 mM dithiothreitol and incubation for 30 minutes at 55°C and then alkylated by the addition of 30 mM iodoacetamide and incubation for 15 minutes at RT in the dark. Phosphoric acid was added to a final concentration of 1.2% and subsequently, samples were diluted 7-fold with binding buffer containing 90% methanol in 100 mM TEAB, pH 7.55. The samples were loaded on the 96-well S-Trap™ plate (Protifi, Fairport, USA), placed on top of a deep-well plate, and centrifuged for 2 min at 1,500 x g at RT. After protein binding, the S-trap™ plate was washed three times by adding 200 μl binding buffer and centrifugation for 2 min at 1,500 x g at RT. A new deep well receiver plate was placed below the 96-well S-Trap™ plate and 50 mM TEAB containing 1 μg trypsin (1/100, w/w) was added for digestion overnight at 37°C. Using centrifugation for 2 min at 1,500 x g, peptides were eluted three times, first with 80 μl 50 mM TEAB, then with 80 μl 0.2% formic acid (FA) in water and finally with 80 μl 0.2% FA in water/acetonitrile (ACN) (50/50, v/v). Eluted peptides were dried completely by vacuum centrifugation. Peptides were dissolved in loading solvent A (0.1% TFA in water/ACN (98:2, v/v)) and desalted on a reversed-phase (RP) C18 OMIX tip (Agilent, Santa Clara, USA). The tip was first washed 3 times with 100 μl pre-wash buffer (0.1% TFA in water/ ACN (20:80, v/v)) and pre-equilibrated 5 times with 100 μl of wash buffer (0.1% TFA in water) before the sample was loaded on the tip. After peptide binding, the tip was washed 3 times with 100 μl of wash buffer and peptides were eluted twice with 100 μl elution buffer (0.1% TFA in water/ACN (40:60, v/v)). The combined elutions were transferred to HPLC inserts and dried in a vacuum concentrator.

### LC-MS/MS analysis

Dried peptides were re-dissolved in 20 μl loading solvent A (0.1% trifluoroacetic acid in water/acetonitrile (ACN) (98:2, v/v)) of which 10 μl was injected for LC-MS/MS analysis on an Ultimate 3000 RSLCnano system in-line connected to a Q Exactive HF mass spectrometer (Thermo Fisher Scientific, Waltham, USA). Trapping was performed at 10 μl/min for 4 min in loading solvent A on a 20 mm trapping column (made in-house, 100 μm internal diameter (I.D.), 5 μm beads, C18 Reprosil-HD, Dr. Maisch, Germany). The peptides were separated on a 250 mm Waters nanoEase M/Z HSS T3 Column, 100Å, 1.8 μm, 75 μm inner diameter (Waters Corporation, Milford, USA) kept at a constant temperature of 45°C. Peptides were eluted by a non-linear gradient starting at 1% MS solvent B reaching 33% MS solvent B (0.1% FA in water/acetonitrile (2:8, v/v)) in 100 min, 55% MS solvent B (0.1% FA in water/acetonitrile (2:8, v/v)) in 135 min, 70% MS solvent B in 145 minutes followed by a 5-minute wash at 99% MS solvent B and re-equilibration with MS solvent A (0.1% FA in water).

The mass spectrometer was operated in data-dependent mode, automatically switching between MS and MS/MS acquisition for the 12 most abundant ion peaks per MS spectrum. Full-scan MS spectra (375-1500 m/z) were acquired at a resolution of 60,000 in the Orbitrap analyzer after accumulation to a target value of 3,000,000. The 12 most intense ions above a threshold value of 15,000 were isolated with a width of 1.5 m/z for fragmentation at a normalized collision energy of 28% after filling the trap at a target value of 100,000 for a maximum of 80 ms. MS/MS spectra (200-2000 m/z) were acquired at a resolution of 15,000 in the Orbitrap analyzer. The polydimethylcyclosiloxane background ion at 445.120028 Da was used for internal calibration (lock mass).

### Bioinformatic analyses

Analysis of the mass spectrometry data for root samples was separately performed in MaxQuant (version 2.0.3.0) with mainly default search settings including a false discovery rate set at 1% on PSM, peptide, and protein level. Spectra were searched against the *Meloidogyne* proteome (TaxId: 189290) (version of January 2022, UniprotKB) containing 72,172 protein sequences and the *Zea Mays* reference proteome (version of January 2022, UP000007305) containing 39,207 protein sequences.

Statistical analyses of significant differences between protein intensities for the datasets obtained from mock-infected and *M. arenaria*-infected plants were done in Perseus software v. 2.0.7.0 (Tyanova et al. 2016). Differentially expressed proteins (DEPs), either with lower and higher abundance were determined using Student’s T-test with p-value < 0.05 and with a cut-off point for log2 fold change (FC) in the range of -0.75 > FC > 0.75.

DEPs with lower and higher abundance were uploaded to the OmicsBox package 3.0 (Bioinformatics Made Easy, BioBam Bioinformatics, March 3, 2019, https://www.biobam.com/omicsbox), and functional annotation was done with Blast2GO tool (Götz et al. 2008) against the *Zea mays* reference proteome and against the InterPro databases (Hunter et al. 2009). Mapping, enrichment analyses, and classification of the gene ontology (GO) terms into functional categories were carried out using default settings in the OmicsBox software with a p-value < 0.05. KAAS, the Kyoto Encyclopedia of Genes and Genomes (KEGG) Automatic Annotation Server (Kanehisa et al. 2023), was used to classify particular proteins into appropriate processes and pathways. KEGGs enrichment analyses were carried out in the OmicsBox software and statistically significant results were determined with p-value < 0.05.

## Results

### Proteome profiling and Differently Expressed Proteins (DEPs) evaluation of healthy and *M. arenaria*-infected roots

During the experiment, a total of 3,743 proteins were identified as belonging to *Z. mays* species, in at least one sample within all tested conditions. In samples collected from healthy plants, at 24 hpi, 1,742 different protein records were identified, while in *M. arenaria* infected plants 1,784 proteins were identified in all three replicates. On the other hand, in samples collected from healthy plants at 3 dpi, 1,805 proteins were identified, while in infected plants 1,529 proteins were observed in all three replicates. Out of all identified proteins 1,258 were common for all experimental conditions (Fig. 1).

**Fig. 1.**
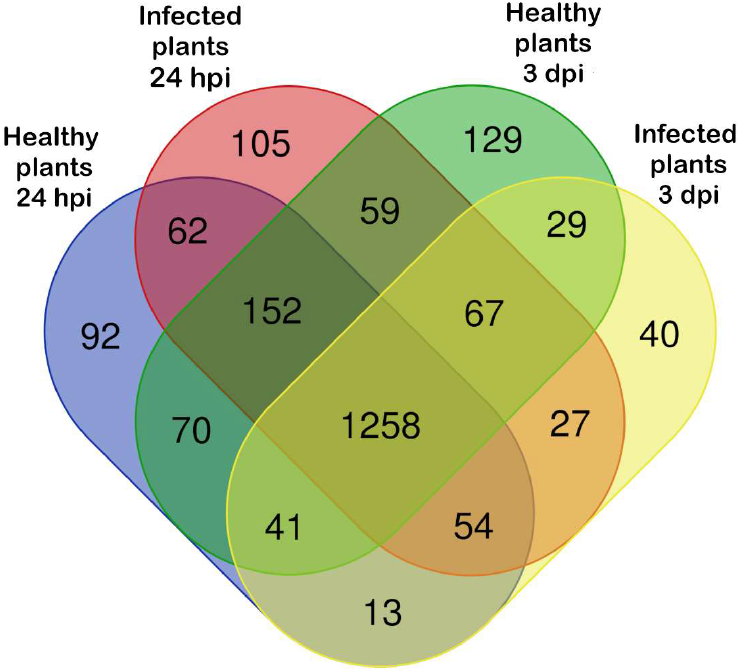
Number of proteins identified in all three replicates of healthy and *Meloidogyne arenaria* infected plants at 24 hours past infection (hpi) and 3 days past infection (dpi). Venn diagram created using the tool available on the website: https://bioinformatics.psb.ugent.be/webtools/Venn

Statistically significant DEPs (p-value < 0.05) were evaluated, by comparing *M. arenaria* infected plants against healthy plants, with the log2 fold change (FC) of -0,75 > FC > 0,75. As a result, 29 downregulated proteins at 24 hpi, 26 proteins at 3 dpi, and one protein common for both time points (an uncharacterized protein with accession number A0A804QSD4) were indicated (Fig. 2A). On the other hand, 41 upregulated proteins were observed at 24 dpi and 96 proteins at 3 dpi, with no proteins common to both time points (Fig. 2B).

**Fig. 2.**
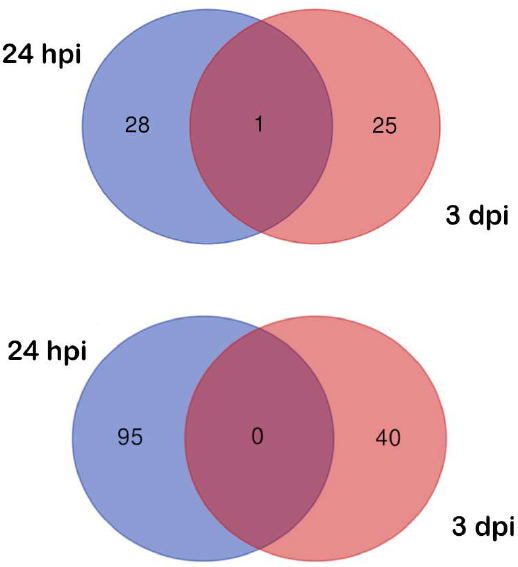
Number of differentially expressed proteins (DEPs) at two time points: 24 hours post inoculation (24 dpi) and 3 days post inoculation (3 dpi). A. Number of downregulated DEPs; B. Number of upregulated DEPs. Venn diagram created using the tool available on the website: https://bioinformatics.psb.ugent.be/webtools/Venn

### GOs enrichment analyses indicate a decrease in the abundance of maize proteins involved in the response to stimulus and removal of superoxide radicals while an increase in proteins involved in cell wall modifications, methylation, phosphorylation, and oxidative stress response

All DEPs, either with decreased or increased abundance, were annotated to GO terms, and enrichment analyses were done. Results of GO term analyses were assigned to biological processes (Table 2), molecular functions (Supplementary Table S1), and cellular components (Supplementary Table S2). Additionally, the individual proteins assigned to the enriched biological processes, with their p-values, were presented in Supplementary Table S3.

The highest number of proteins with decreased abundance at 24 hpi was assigned to three biological processes: sulfur compound metabolic process (3 proteins), aerobic respiration (3 proteins), and electron transport chain (3 proteins) while at 3 dpi there were two other processes: cellular response to stimulus (6 proteins) and response to heat (3 proteins) (Fig. 3A). On the other hand, from biological processes enriched by proteins with increased abundance at 24 hpi, five were the most numerous represented: response to oxidative stress (7 proteins), cellular oxidant detoxification (7 proteins), protein phosphorylation (5 proteins), hydrogen peroxide catabolic process (5 proteins), and defense response (4 proteins). Moreover, at 3 dpi, five other processes were significantly overrepresented: methylation (5 proteins), phenylpropanoid metabolic process (5 proteins), fatty acid catabolic process (3 proteins), one carbon metabolic process (3 proteins), and S-adenosylmethionine biosynthetic process (3 proteins) (Fig. 3B).

**Fig. 3.**
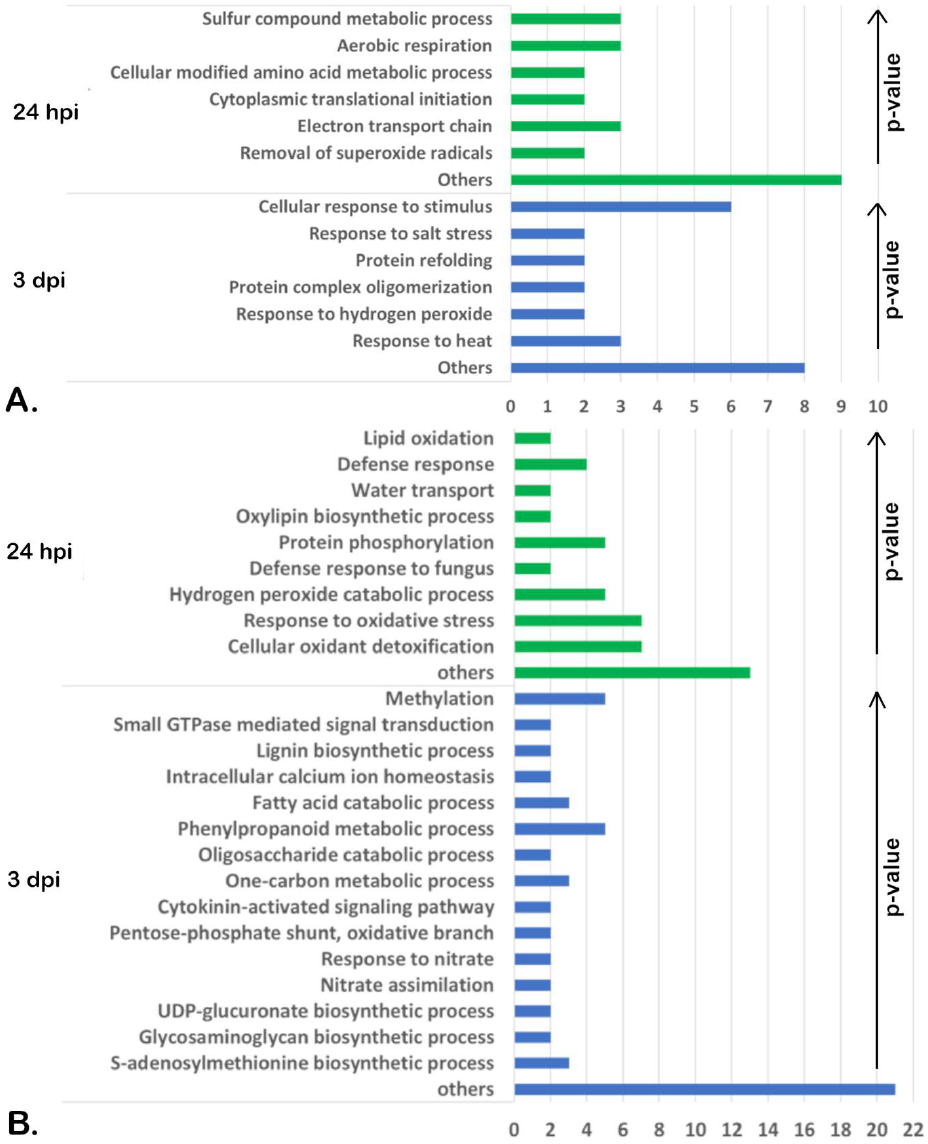
Results of enrichment analysis of significantly differentially expressed proteins (DEPs) assigned to selected biological processes at 24 hours post inoculation (hpi) and 3 days post inoculation (dpi) in *Zea mays* after *Meloidogyne arenaria* infection. A. Proteins with decreased accumulation level; B. Proteins with increased accumulation level.

### KEGGs enrichment analyses indicate changes in maize proteins related to lipid metabolism, biosynthesis of secondary metabolites, and signal transduction after *M. arenaria* infection

Down- and upregulated proteins from samples collected at 24 hpi and 3 dpi were classified into appropriate KEGG processes and pathways followed by enrichment analyses (Table 3). Additionally, the individual proteins assigned to the enriched KEGG processes and pathways were presented in Supplementary Table S4.

**Table 1.**
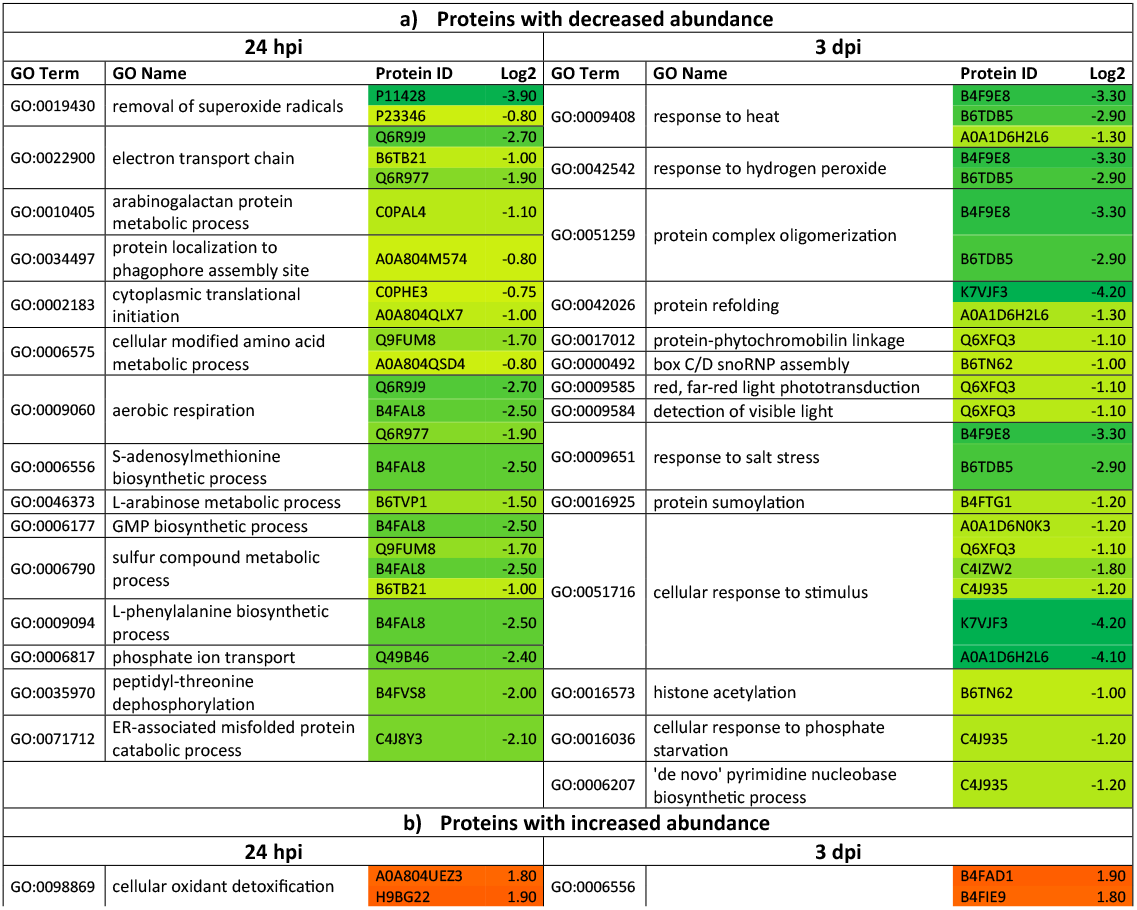
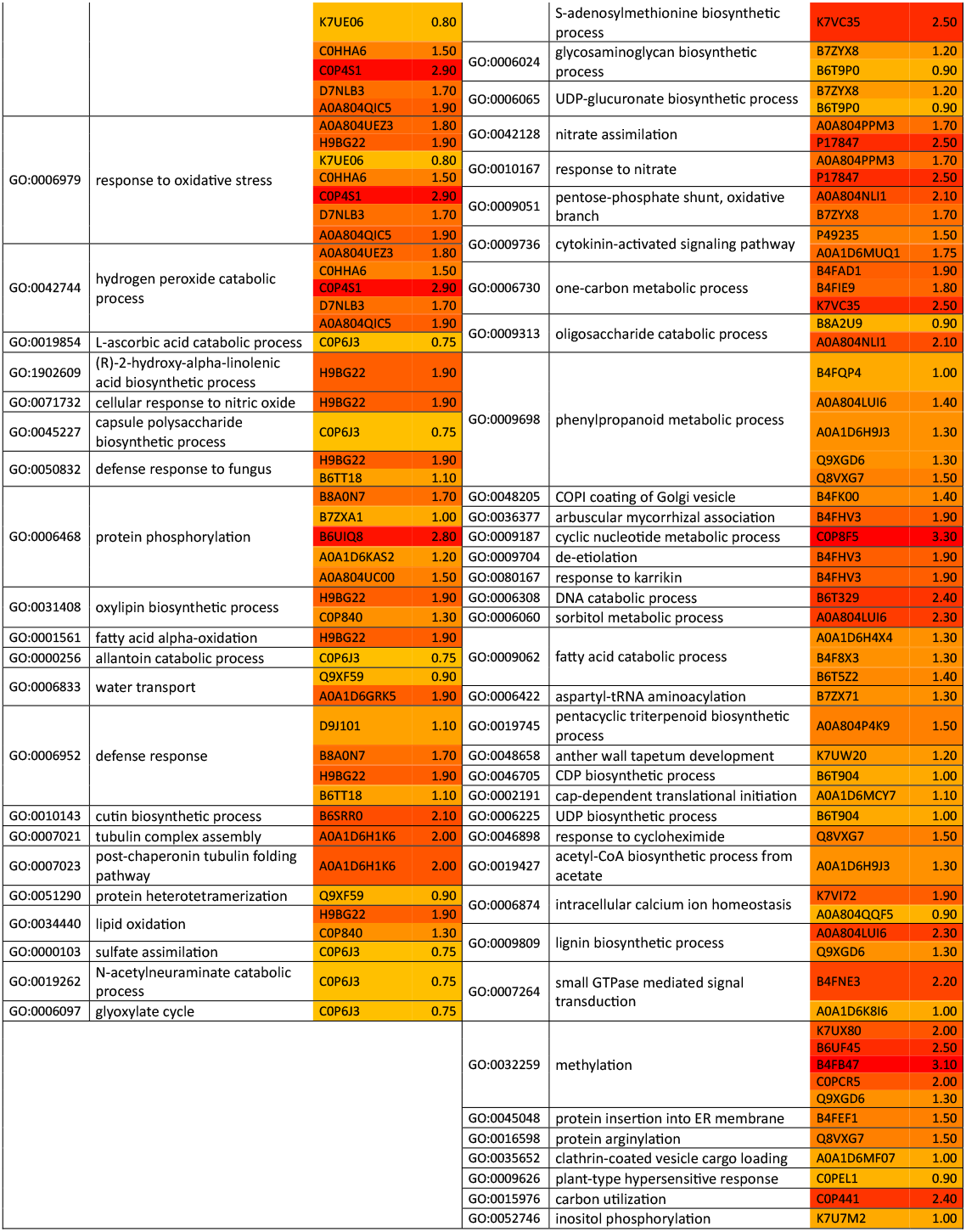
Biological processes enriched at 24 hours post inoculation (hpi) and 3 days post inoculation (dpi) with differently expressed proteins (DEPs) either with a) lower and b) higher abundance with Log2 fold change for proteins assigned to each process. GO terms p-values provided in Suppl. Table S3.

**Table 3.** Proteins with decreased (red arrows)- and increased (green arrows) accumulation levels, assessed do each Kyoto Encyclopedia of Genes and Genomes (KEGGs) process and pathway at 24 hours post inoculation (hpi) and 3 days post inoculation (dpi)

| Pathway group | Pathway subgroup | Pathway | 24 hpi |  | 3 dpi |  |
| --- | --- | --- | --- | --- | --- | --- |
|  |  |  |  | p-value |  | p-value |
| Metabolism | Global and overview maps | Biosynthesis of nucleotide sugars |  |  | ↑4 | 4.98E-02 |
|  |  | Biosynthesis of cofactors |  |  | ↑8 | 2.34E-02 |
|  | Carbohydrate metabolism | Pentose and glucuronate interconversions |  |  | ↑4 | 1.90E-03 |
|  | Energy metabolism | Nitrogen metabolism |  |  | ↑4 | 1.90E-03 |
|  | Lipid metabolism | alpha-Linolenic acid metabolism | ↑4 | 1.71E-04 |  |  |
|  |  | Steroid biosynthesis |  |  | ↑2 | 3.59E-02 |
|  | Amino acid metabolism | Cysteine and methionine metabolism |  |  | ↑5 | 2.89E-02 |
|  | Glycan biosynthesis and metabolism | O-Antigen nucleotide sugar biosynthesis |  |  | ↑2 | 1.98E-02 |
|  | Biosynthesis of other secondary metabolites | Phenylpropanoid biosynthesis | ↑6 | 2.86E-04 | ↑8 | 7.47E-03 |
|  |  | Stilbenoid, diarylheptanoid, and gingerol biosynthesis |  |  | ↑3 | 3.28E-02 |
|  |  | Flavonoid biosynthesis |  |  | ↑3 | 3.53E-02 |
|  |  | Neomycin, kanamycin and gentamicin biosynthesis | ↑1 | 4.72E-02 |  |  |
|  |  | Prodigiosin biosynthesis |  |  | ↓1 | 4.49E-02 |
|  |  | Biosynthesis of various secondary metabolites |  |  | ↑5 | 1.50E-05 |
|  | Folding, sorting and degradation | Protein processing in endoplasmic reticulum |  |  | ↓3 | 2.69E-02 |
| Environmental Information Processing | Signal transduction | Wnt signaling pathway |  |  | ↑3 | 1.05E-02 |
|  |  | MAPK signaling pathway | ↑2 | 4.36E-02 |  |  |
|  |  | MAPK signaling pathway plant |  |  | ↓2 | 3.06E-02 |
|  |  | NF-kappa B signaling pathway | ↑2 | 1.53E-02 |  |  |
|  |  | Plant hormone signal transduction |  |  | ↓2 | 2.03E-02 |
|  |  | Peroxisome | ↓2 | 3.80E-02 |  |  |
|  | Cellular community | Adherens junction |  |  | ↑2 | 2.74E-02 |
| Organismal systems | Immune system | Toll-like receptor signaling pathway | ↑2 | 1.97E-02 |  |  |
|  |  | Toll and lmd signaling pathway | ↑2 | 1.97E-02 |  |  |
|  |  | Leukocyte transendothelial migration |  |  | ↑1 | 4.02E-02 |
|  | Nervous system | Retrograde endocannabinoid signaling | ↓2 | 1.43E-02 |  |  |
|  |  | Neurotrophin signaling pathway | ↑2 | 3.28E-02 |  |  |
|  | Development and regeneration | Axon guidance |  |  | ↑2 | 2.35E-02 |
|  | Aging | Longevity regulating pathway – multiple species | ↓2 | 1.22E-02 | ↓2 | 1.06E-02 |

### thway 24 hpi 3 dpi

As a result of KEGG enrichment analyses of proteins with decreased abundance, it was observed that they were assigned to the longevity-regulating pathway both at 24 hpi and 3 dpi (2 proteins). It is also worth noting that proteins with decreased accumulation levels at 24 hpi were involved in signal transduction in peroxisome (2 proteins), but at 3 dpi in MAPK signaling pathway (2 proteins) and plant hormone signal transduction (2 proteins) (Fig. 4A). The KEGGs analyses of proteins with the higher abundance revealed that the most significantly enriched KEGG pathway in both time points was the phenylpropanoid biosynthesis pathway (6 proteins at 24 hpi and 8 proteins at 3 dpi). Other important pathways overrepresented at 24 hpi were alpha-linolenic acid metabolism (4 proteins) and MAPK signaling pathway (2 proteins), while at 3 dpi, the biosynthesis of cofactors (8 proteins), cysteine and methionine metabolism (5 proteins), nitrogen metabolism (4 proteins) and flavonoid biosynthesis (3 proteins) are worth mentioning (Fig. 4B).

**Fig. 4.**
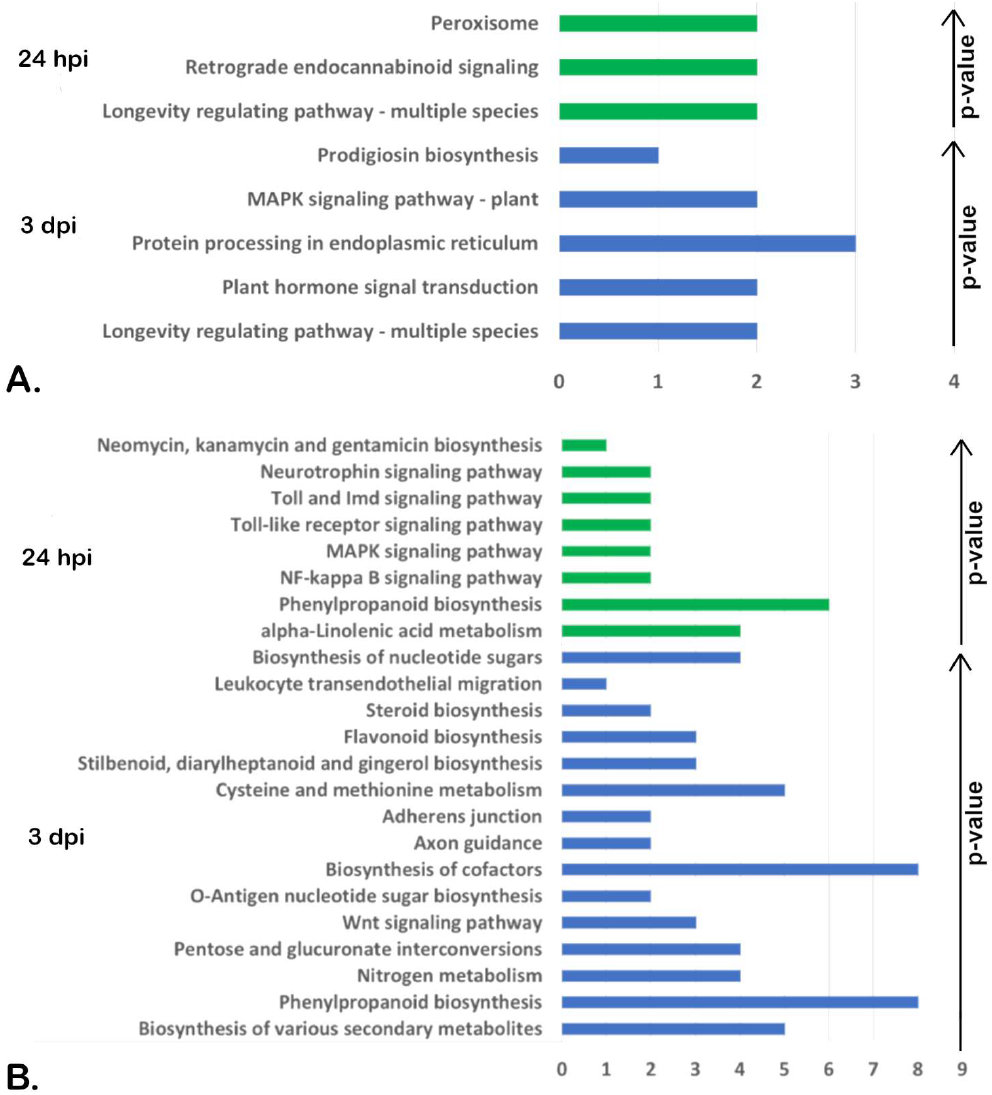
Results of enrichment analysis of differentially expressed proteins (DEPs) assigned to Kyoto Encyclopedia of Genes and Genomes (KEGGs) processes and pathways at 24 hours post inoculation (hpi) and 3 days post inoculation (dpi) in *Zea mays* after *Meloidoryne arenaria* infection. A. Proteins with decreased accumulation level; B. Proteins with increased accumulation level.

## Discussion

Our aim was to describe global changes in the root proteome of a susceptible maize variety during the early stage of *M. arenaria* infection. The profiling of the proteome and transcriptome of RKN-infected monocotyledonous hosts has been the subject of only a few studies, none of which involved maize. In our previous study, we observed some changes in susceptible varieties of maize compared to tolerant ones, but they were verified on the transcriptomic level for only a few proteins, related mainly to plant defense response, cell wall modification, and regulation of genes expression (Przybylska et al. 2018; Przybylska and Spychalski 2021). Recently, we also identified a potential *M. arenaria* effector MaMsp4, and found out that it interacts with proteins associated with polyamines and phosphatidylethanolamine biosynthesis, and with those involved in cell wall modifications during compatible interactions with maize (Przybylska et al. 2023).

The first layer of plant defense against pests is the cell wall, composed mainly of cellulose, hemicelluloses, and pectins, constituting a complicated, three-dimensional structure (Keegstra 2010). The role of proteins related to cell wall modifications in plant–RKNs interactions was the subject of our previous studies conducted in the *M. arenaria* – maize pathosystem (Przybylska and Spychalski 2021; Przybylska et al. 2023). We observed significant upregulation of genes encoding polygalacturonase, beta-galactosidase, and pectinesterase at 24 hpi in samples from roots from a susceptible variety of maize. Moreover, genes encoding such proteins as endoglucanase, xyloglucan endotransglucosylase, and cellulose synthase were upregulated in a susceptible variety of tomatoes in the early stage of *M. incognita* infection (Shukla et al. 2018). Induction of genes related to cell wall modifications was also observed during *Solanum torvum*’s response to *M. arenaria* infection (Sato et al. 2021). These data overlap with results described in this study, where changes in the accumulation level of proteins related to cell wall modifications were observed at both time points. In enriched GO terms assigned to biological processes, the higher abundance of proteins assigned to the cutin biosynthetic process was observed at 24 hpi and those related to the lignin biosynthetic process were indicated at 3 dpi (Table 2), while in GO terms assigned to cellular component, the plant-type cell wall was overrepresented at 24 hpi (Supplementary Table S2). All these results confirm that the remodeling of the cell wall structure is essential for the successful penetration of RKN into the plant tissue and the use of its resources.

Plant defense response is related to many mechanisms and involves among others ROS burst or induction of JA- and SA-related pathways (Manosalva et al. 2015). Antioxidants, such as ascorbate, glutathione, tocopherols, flavonoids, superoxide dismutase, catalase, glutathione S-transferase or peroxidase take part in scavenging of ROS and protect the plants from oxidative damage (Hasanuzzaman and Fujita 2022; Frąckowiak et al. 2022). On the other hand, lipids and fatty acids as well as products generated during fatty acid metabolism play an important role in signaling during plant defense (Lim et al. 2017). In this study, we observed significant changes in processes and pathways related to plant defense response in maize at 24 hpi. From GO enriched terms, the increase in accumulation level was observed for proteins assigned to, among others, defense response, defense response to fungus, response to oxidative stress, hydrogen peroxide catabolic process, oxylipin biosynthetic process, and lipid oxidation (Table 2), while in KEGGs results those proteins were associated with alpha-linolenic acid metabolism (Table 3). However, proteins assigned to flavonoid biosynthesis were overrepresented at 3 dpi. The major proteins annotated to defense response, response to oxidative stress, and lipid oxidation processes were peroxidases, alpha-dioxygenase, allene oxide synthesis, lipoxygenase, and benzoate O-methyltransferase (Supplementary Tables S3 and S4). Interestingly, the expression of genes encoding class III peroxidases was induced at 24 hpi, followed by the induction of the expression of genes involved in defense response at 3 dpi during compatible interactions between *M. arenaria* and *S. torvum* (Sato et al. 2021), which partially overlaps with results observed in this study. On the other hand, we previously reported downregulation of gene encoding peroxidase in all analyzed time points (24 hpi, 3 dpi, and 7 dpi) in maize response to *M. arenaria* infection. Moreover, we observed upregulation lipoxygenase coding gene only at 7 dpi (Przybylska et al. 2018), in contrast to the results obtained in this study, where a higher accumulation of lipooxygenase was observed in the very early stage of infection. Significant upregulation of proteins related to response to oxidative stress and fatty acid metabolism only at 24 hpi suggests that plant defense mechanism is induced in the very early stage of infection but this response might be suppressed by the RKN in the later stages.

Another process, reported to play an important role in plant defense response in many pathosystems is phenylpropanoid biosynthesis and metabolism. The phenylpropanoid pathway is related to the biogenesis of several compounds such as flavonoids, monolignols, phenolic acids, stilbenes, phytoalexins, and coumarins, which are directly and indirectly involved in plant development and disease response. Moreover, this process entails cell wall lignification, which is one of the first layers of plant defense (Yadav et al. 2020). Phenylpropanoid metabolism was particularly enriched in the results of proteome and transcriptome analysis of the soybean response to *M. incognita* infection (Arraes et al. 2022) which is consistent with GO terms from this study, in which we observed an increase in the abundance of proteins related to this process at 3 dpi (Table 2). Likewise, phenylpropanoid biosynthesis was upregulated during tomato response to *M. incognita* infection (Shukla et al. 2018). It overlaps with KEGGs results obtained in this study, where a higher abundance of proteins assigned to phenylpropanoid biosynthesis was observed at both analyzed time points (Table 3). Noteworthy, the *M. javanica* effector protein, Mj-FAR-1, was shown to regulate the expression of phenylpropanoid-related genes during the infection of tomato (Iberkleid et al. 2015). One of the key enzymes annotated to the phenylpropanoid pathway is phenylalanine/tyrosine ammonia-lyase (PAL), which is involved in abiotic and biotic stress responses (Barros and Dixon 2020).

The PAL-mediated SA biosynthesis was reported to contribute to wheat resistance to cereal cyst nematode (Zhang et al. 2021). In our study, we observed a significant increase in the accumulation level of this protein at 3 dpi. Results obtained in this and other studies altogether suggest that induction of processes and pathways related to phenylpropanoid are fundamental in the host response to RKN infection. A higher abundance of proteins related to these processes occurs at both analyzed time points, in contrast to proteins related to oxidative stress and fatty acid metabolism, overrepresented only at 24 hpi.

Among other proteins related to plant defense response, it is worth mentioning an S-adenosylmethionine, which is a substrate for S-adenosylmethionine decarboxylase in the polyamines biosynthesis pathway. The role of polyamines was investigated in the regulation of cell division, root formation, flower and fruit development, cell wall generation, xylem differentiation, leaf senescence, and programmed cell death, but also in plant defense signaling (Asija et al. 2022). In our previous study, we reported, that S-adenosylmethionine decarboxylase is a target for *M. arenaria* potential effector protein in maize, MaMsp4 (Przybylska et al. 2023). Moreover, we observed the downregulation of the gene encoding S-adenosylmethionine decarboxylase 2 in the early stage of infection (24 hpi) during compatible interactions, which may suggest inhibition of polyamines biosynthesis. On the other hand, in this study, we observed that the accumulation level of three S-adenosylmethionine synthases was significantly increased at 3 dpi. They were assigned to the cysteine and methionine metabolism, biosynthesis of cofactors, and biosynthesis of various secondary metabolites in KEGG results (Supplementary Table S4), and to the S-adenosylmethionine biosynthetic process (Table 3) and one-carbon metabolic biological process (Supplementary Table S3) in GO terms. Moreover, S-adenosylmethionine synthase was also significantly overrepresented in another monocotyledonous host of RKN - banana at 60 dpi after *M. incognita* infection (Al-Idrus et al. 2017). However, we observed a decrease in the accumulation level of one protein assigned to the S-adenosylmethionine biosynthetic process at 24 hpi (Table 3), as well as protein with S-methylmethionine-homocysteine S-methyltransferase activity (Supplementary Table S1). Interestingly, we have not observed significant changes in other processes and pathways related to polyamines biosynthesis in the analyzed time points.

Other important pathways and processes are those related to signal transduction. MAPK signal transduction pathway is a signal cascade consisting of receptors and sensors that transduce extracellular stimuli into intracellular responses in cells (Zhang and Klessig 2001). On the other hand, cytokinin signal transduction occurs through a multistep phosphorelay that incorporates receptors with histidine–kinase activity, phosphotransfer proteins, or type-B response regulators (Zubo and Schaller 2020). Here, we observed a higher abundance of proteins assigned to the MAPK signaling pathway, NF-kappa B signaling pathway, and Toll-like receptor signaling pathways at 24 hpi, while proteins related to the Wnt signaling pathway at 3 dpi in KEGGs results (Table 3). However, a decrease in the accumulation level of proteins assigned to the MAPK signaling pathway in plant and plant hormone signal transduction pathway was observed at 3 dpi. Interestingly, PR-1 protein was associated with both these pathways (Supplementary Table S4). Moreover, the lower abundance of PR-1 protein was also observed in compatible interactions between banana and *M. incognita* (Al-Idrus et al. 2017) as well as downregulation of its transcript level in tomato after *M. incognita* infection (Shukla et al. 2018). Noteworthy, the MAPK signaling pathway was reported to be overrepresented during rice response to *M. incognita* infection (Zhou et al. 2020). Signal transduction pathways were also enriched during the mulberry response to *M. enterolobii* infection (Shao et al. 2021). Interestingly, in GO term results, proteins assigned to cytokinin-activated signaling pathway were on higher accumulation level at 3 dpi (Table 2). It is consistent with data obtained by Castaneda et al. (2017), where overexpression of transcript of gene encoding protein related to cytokinin-activated signaling pathway was also observed at the same time point during the banana response to *M. incognita* infection. Considering these results together, it can be observed that MAPK signal transduction is induced at a very early stage (24 hpi), but then partially suppressed, while cytokinin-activated signaling pathway, related to phosphorylation is activated later (3 dpi).

## Conclusions

In this study, we observed significant changes in a number of processes and pathways in maize at an early stage of *M. arenaria* infection. In addition to the role of processes such as cell wall modifications, plant defense response mechanisms, phenylpropanoid metabolism, and signal transduction pathways, it is worth mentioning the process of S-adenosylmethionine biosynthesis, significantly upregulated at 3 dpi. Consistent with this, as well as with our previous study, S-adenosylmethionine might play an important role in compatible interactions between maize and *M. arenaria*.

## Acknowledgment

Proteins samples preparation and LC-MS/MS LFQ analyses were done by Sara Dufour, Delphi Van Haver, and Francis Impens in VIB-UGent Center for Medical Biotechnology (Ghent, Belgium) within the Epic-XS project, as a part of Horizon2020. This study was also supported by a grant from the Polish National Science Center [grant number 2014/13/N/NZ9/00703] and by a project from the Ministry of Education and Science [BIOTECH-01].

## Supplementary tables

**Supplementary Table SF1**. Results of enrichment analysis of proteins assigned to particular molecular functions at 24 hpi and 3 dpi with differently expressed proteins (DEPs) either with lower and higher abundance.

**Supplementary Table S2**. Results of enrichment analysis of proteins assigned to particular cellular components at 24 hpi and 3 dpi with differently expressed proteins (DEPs) either with lower and higher abundance.

**Supplementary Table S3**. Individual proteins with higher and lower abundance, assessed to biological processes enriched at 24 hpi and 3 dpi.

**Supplementary Table S4**. Individual proteins with lower and higher abundance, assessed to each KEGG process and pathway at 24 hpi and 3 dpi.

