## Supplementary Tables for "Profiling of *Zea mays* L. proteome at early stages of compatible interactions with *Meloidogyne arenaria* indicates changes in signaling, oxidative stress responses, and S-adenosylmethionine biosynthesis"

**Supplementary Table S1**. Results of enrichment analysis of proteins assigned to particular molecular functions at 24 hpi and 3 dpi with differently expressed proteins (DEPs) either with lower and higher abundance.

| **Molecular functions** | | | |
|---|---|---|---|
| **GO Term** | **GO Name** | **P-value** | **Nr test** |
| **Proteins with lower abundance** | | | |
| **24 hpi** | | | |
| GO:0004784 | superoxide dismutase activity | 4.43E-04 | 2 |
| GO:0008137 | NADH dehydrogenase (ubiquinone) activity | 1.62E-03 | 2 |
| GO:0018549 | methanethiol oxidase activity | 4.71E-03 | 1 |
| GO:0061627 | S-methylmethionine-homocysteine S-methyltransferase activity | 4.71E-03 | 1 |
| GO:0008430 | selenium binding | 4.71E-03 | 1 |
| GO:0016844 | strictosidine synthase activity | 4.71E-03 | 1 |
| GO:0080041 | ADP-ribose pyrophosphohydrolase activity | 4.71E-03 | 1 |
| GO:0080042 | ADP-glucose pyrophosphohydrolase activity | 4.71E-03 | 1 |
| GO:0005507 | copper ion binding | 5.10E-03 | 2 |
| GO:0017018 | myosin phosphatase activity | 1.36E-02 | 2 |
| GO:0004477 | methenyltetrahydrofolate cyclohydrolase activity | 1.41E-02 | 1 |
| GO:0004488 | methylenetetrahydrofolate dehydrogenase (NADP+) activity | 1.41E-02 | 1 |
| GO:0042301 | phosphate ion binding | 1.87E-02 | 1 |
| GO:0140658 | ATP-dependent chromatin remodeler activity | 1.87E-02 | 1 |
| GO:0047938 | glucose-6-phosphate 1-epimerase activity | 1.87E-02 | 1 |
| GO:0046524 | sucrose-phosphate synthase activity | 1.87E-02 | 1 |
| GO:0052692 | raffinose alpha-galactosidase activity | 2.80E-02 | 1 |
| GO:0004365 | glyceraldehyde-3-phosphate dehydrogenase (NAD+) (phosphorylating) activity | 3.25E-02 | 1 |
| GO:0005315 | inorganic phosphate transmembrane transporter activity | 4.17E-02 | 1 |
| GO:0016157 | sucrose synthase activity | 4.17E-02 | 1 |
| GO:0070569 | uridylyltransferase activity | 4.62E-02 | 1 |
| **3 dpi** | | | |
| GO:0043621 | protein self-association | 3.40E-03 | 2 |
| GO:0051082 | unfolded protein binding | 3.64E-03 | 3 |
| GO:0030246 | carbohydrate binding | 7.06E-03 | 3 |

| | | | |
|---|---|---|---|
| GO:0019948 | SUMO activating enzyme activity | 8.40E-03 | 1 |
| GO:0004070 | aspartate carbamoyltransferase activity | 8.40E-03 | 1 |
| GO:0004477 | methenyltetrahydrofolate cyclohydrolase activity | 1.26E-02 | 1 |
| GO:0004488 | methylenetetrahydrofolate dehydrogenase (NADP+) activity | 1.26E-02 | 1 |
| GO:0016877 | ligase activity, forming carbon-sulfur bonds | 1.84E-02 | 2 |
| GO:0009881 | photoreceptor activity | 2.91E-02 | 1 |
| GO:0000155 | phosphorelay sensor kinase activity | 2.91E-02 | 1 |
| GO:0016597 | amino acid binding | 4.13E-02 | 1 |
| **Proteins with higher abundance** | | | |
| **24 hpi** | | | |
| GO:0020037 | heme binding | 1.95E-06 | 9 |
| GO:0140825 | lactoperoxidase activity | 7.60E-05 | 5 |
| GO:0043167 | ion binding | 3.29E-03 | 23 |
| GO:0004674 | protein serine/threonine kinase activity | 5.56E-03 | 5 |
| GO:0016702 | oxidoreductase activity, acting on single donors with incorporation of molecular oxygen, incorporation of two atoms of oxygen | 6.32E-03 | 2 |
| GO:0080150 | S-adenosyl-L-methionine:benzoic acid carboxyl methyl transferase activity | 6.73E-03 | 1 |
| GO:0043014 | alpha-tubulin binding | 6.73E-03 | 1 |
| GO:0015250 | water channel activity | 7.03E-03 | 2 |
| GO:0004020 | adenylylsulfate kinase activity | 1.34E-02 | 1 |
| GO:0016162 | cellulose 1,4-beta-cellobiosidase activity | 3.32E-02 | 1 |
| GO:0090447 | glycerol-3-phosphate 2-O-acyltransferase activity | 2.67E-02 | 1 |
| GO:0005536 | glucose binding | 4.62E-02 | 1 |
| GO:0102726 | DIMBOA glucoside beta-D-glucosidase activity | 4.62E-02 | 1 |
| GO:0004340 | glucokinase activity | 4.62E-02 | 1 |
| GO:0019199 | transmembrane receptor protein kinase activity | 1.11E-02 | 2 |
| GO:0004634 | phosphopyruvate hydratase activity | 3.32E-02 | 1 |
| GO:0008115 | sarcosine oxidase activity | 2.67E-02 | 1 |
| GO:0004781 | sulfate adenylyltransferase (ATP) activity | 1.34E-02 | 1 |
| GO:0009978 | allene oxide synthase activity | 1.34E-02 | 1 |
| **3 dpi** | | | |
| GO:0004478 | methionine adenosyltransferase activity | 1.47E-05 | 3 |

| | | | |
|---|---|---|---|
| GO:0048307 | ferredoxin-nitrite reductase activity | 2.42E-04 | 2 |
| GO:0003979 | UDP-glucose 6-dehydrogenase activity | 2.42E-04 | 2 |
| GO:0016162 | cellulose 1,4-beta-cellobiosidase activity | 2.35E-03 | 2 |
| GO:0102726 | DIMBOA glucoside beta-D-glucosidase activity | 4.84E-03 | 2 |
| GO:0015928 | fucosidase activity | 6.38E-03 | 2 |
| GO:0102483 | scopolin beta-glucosidase activity | 1.22E-02 | 2 |
| GO:0031491 | nucleosome binding | 1.44E-02 | 2 |
| GO:0047172 | shikimate O-hydroxycinnamoyltransferase activity | 1.57E-02 | 1 |
| GO:0004113 | 2',3'-cyclic-nucleotide 3'-phosphodiesterase activity | 1.57E-02 | 1 |
| GO:0016508 | long-chain-enoyl-CoA hydratase activity | 1.57E-02 | 1 |
| GO:0016594 | glycine binding | 1.57E-02 | 1 |
| GO:0009010 | sorbitol-6-phosphate 2-dehydrogenase activity | 1.57E-02 | 1 |
| GO:0004375 | glycine dehydrogenase (decarboxylating) activity | 1.57E-02 | 1 |
| GO:0045430 | chalcone isomerase activity | 1.57E-02 | 1 |
| GO:0004612 | phosphoenolpyruvate carboxykinase (ATP) activity | 1.57E-02 | 1 |
| GO:0046526 | D-xylulose reductase activity | 1.57E-02 | 1 |
| GO:0003939 | L-iditol 2-dehydrogenase activity | 1.57E-02 | 1 |
| GO:0000014 | single-stranded DNA endodeoxyribonuclease activity | 1.57E-02 | 1 |
| GO:0008670 | 2,4-dienoyl-CoA reductase (NADPH) activity | 1.57E-02 | 1 |
| GO:0050255 | ribitol 2-dehydrogenase activity | 1.57E-02 | 1 |
| GO:0015923 | mannosidase activity | 1.69E-02 | 2 |
| GO:0070403 | NAD+ binding | 1.95E-02 | 2 |
| GO:0097599 | xylanase activity | 2.23E-02 | 2 |
| GO:0050661 | NADP binding | 3.10E-02 | 3 |
| GO:0035299 | inositol pentakisphosphate 2-kinase activity | 3.11E-02 | 1 |
| GO:0098808 | mRNA cap binding | 3.11E-02 | 1 |
| GO:0042409 | caffeoyl-CoA O-methyltransferase activity | 3.11E-02 | 1 |
| GO:0016401 | palmitoyl-CoA oxidase activity | 3.11E-02 | 1 |
| GO:0004324 | ferredoxin-NADP+ reductase activity | 3.11E-02 | 1 |
| GO:0016871 | cycloartenol synthase activity | 3.11E-02 | 1 |
| GO:0047918 | GDP-mannose 3,5-epimerase activity | 3.11E-02 | 1 |
| GO:0004815 | aspartate-tRNA ligase activity | 3.11E-02 | 1 |

| | | | |
|---|---|---|---|
| GO:0004615 | phosphomannomutase activity | 3.11E-02 | 1 |
| GO:0003987 | acetate-CoA ligase activity | 3.11E-02 | 1 |
| GO:0052747 | sinapyl alcohol dehydrogenase activity | 3.11E-02 | 1 |
| GO:0052883 | tyrosine ammonia-lyase activity | 3.11E-02 | 1 |
| GO:0005085 | guanyl-nucleotide exchange factor activity | 3.49E-02 | 2 |
| GO:0015926 | glucosidase activity | 4.08E-02 | 3 |
| GO:0036430 | CMP kinase activity | 4.62E-02 | 1 |
| GO:0036431 | dCMP kinase activity | 4.62E-02 | 1 |
| GO:0004591 | oxoglutarate dehydrogenase (succinyl-transferring) activity | 4.62E-02 | 1 |
| GO:0004575 | sucrose alpha-glucosidase activity | 4.62E-02 | 1 |
| GO:0004838 | L-tyrosine:2-oxoglutarate aminotransferase activity | 4.62E-02 | 1 |
| GO:0004616 | phosphogluconate dehydrogenase (decarboxylating) activity | 4.62E-02 | 1 |
| GO:0033926 | glycopeptide alpha-N-acetylgalactosaminidase activity | 4.62E-02 | 1 |
| GO:0033862 | UMP kinase activity | 4.62E-02 | 1 |
| GO:0008692 | 3-hydroxybutyryl-CoA epimerase activity | 4.62E-02 | 1 |
| GO:0015662 | P-type ion transporter activity | 4.96E-02 | 2 |

**Supplementary Table S2.** Results of enrichment analysis of proteins assigned to particular cellular components at 24 hpi and 3 dpi with differently expressed proteins (DEPs) either with lower and higher abundance.

| **Cellular components** | | | |
|---|---|---|---|
| **GO Term** | **GO Name** | **P-value** | **Nr test** |
| **Proteins with lower abundance** | | | |
| **24 hpi** | | | |
| GO:0005758 | mitochondrial intermembrane space | 1.41E-02 | 1 |
| GO:1990072 | TRAPPIII protein complex | 1.41E-02 | 1 |
| GO:0000407 | phagophore assembly site | 2.80E-02 | 1 |
| GO:0034098 | VCP-NPL4-UFD1 AAA ATPase complex | 4.17E-02 | 1 |
| GO:0010494 | cytoplasmic stress granule | 4.62E-02 | 1 |
| **3 dpi** | | | |
| GO:0031510 | SUMO activating enzyme complex | 8.40E-03 | 1 |
| GO:0097255 | R2TP complex | 1.67E-02 | 1 |

| | | | |
|---|---|---|---|
| GO:0000812 | Swr1 complex | 2.09E-02 | 1 |
| GO:0035267 | NuA4 histone acetyltransferase complex | 2.50E-02 | 1 |
| GO:0031011 | Ino80 complex | 2.91E-02 | 1 |
| **Proteins with higher abundance** | | | |
| **24 hpi** | | | |
| GO:0005911 | cell-cell junction | 1.67E-04 | 5 |
| GO:0009505 | plant-type cell wall | 1.80E-04 | 4 |
| GO:0012511 | monolayer-surrounded lipid storage body | 6.73E-03 | 1 |
| GO:0005576 | extracellular region | 2.91E-02 | 5 |
| GO:0034515 | proteasome storage granule | 3.97E-02 | 1 |
| GO:0000015 | phosphopyruvate hydratase complex | 3.32E-02 | 1 |
| **3 dpi** | | | |
| GO:0009507 | chloroplast | 2.98E-02 | 11 |
| GO:0035101 | FACT complex | 3.11E-02 | 1 |
| GO:0005829 | cytosol | 3.70E-02 | 18 |
| GO:0005960 | glycine cleavage complex | 4.62E-02 | 1 |

**Supplementary Table S3**. Individual proteins with higher and lower abundance, assessed to biological processes enriched at 24 hpi and 3 dpi.

| Proteins with lower abundance | | | | | | |
|---|---|---|---|---|---|---|
| 24 hpi | | | | | | |
| **GO Term** | **GO Name** | **P-value** | **Nr test** | **Assigned proteins names** | **Protein ID** | **Log2 Fold Change** |
| GO:0019430 | removal of superoxide radicals | 9.42E-04 | 2 | Superoxide dismutase [Cu-Zn] 2 | P11428 | -3,9 |
| | | | | Superoxide dismutase [Cu-Zn] 4AP | P23346 | -0,8 |
| GO:0022900 | electron transport chain | 7.15E-03 | 3 | NADH dehydrogenase subunit 2 | Q6R9J9 | -2,7 |
| | | | | Anamorsin homolog | B6TB21 | -1,0 |
| | | | | NADH-ubiquinone oxidoreductase chain 1 | Q6R977 | -1,9 |
| GO:0010405 | arabinogalactan protein metabolic process | 9.40E-03 | 1 | Alpha-galactosidase | C0PAL4 | -1,1 |
| GO:0034497 | protein localization to phagophore assembly site | 9.40E-03 | 1 | Uncharacterized protein A0A804M574 | A0A804M574 | -0,8 |
| GO:0002183 | cytoplasmic translational initiation | 1.36E-02 | 2 | Eucaryotic initiation factor4 | C0PHE3 | -0,75 |
| | | | | Uncharacterized protein A0A804QLX7 | A0A804QLX7 | -1,0 |
| GO:0006575 | cellular modified amino acid metabolic process | 1.79E-02 | 2 | Homocysteine S-methyltransferase 3 | Q9FUM8 | -1,7 |
| | | | | Uncharacterized protein A0A804QSD4 | A0A804QSD4 | -0,8 |
| GO:0009060 | aerobic respiration | 2.73E-02 | 3 | NADH dehydrogenase subunit 2 | Q6R9J9 | -2,7 |
| | | | | Methanethiol oxidase | B4FAL8 | -2,5 |
| | | | | NADH-ubiquinone oxidoreductase chain 1 | Q6R977 | -1,9 |
| GO:0006556 | S-adenosylmethionine biosynthetic process | 2.80E-02 | 1 | Methanethiol oxidase | B4FAL8 | -2,5 |
| GO:0046373 | L-arabinose metabolic process | 3.25E-02 | 1 | Uncharacterized protein B6TVP1 | B6TVP1 | -1,5 |
| GO:0006177 | GMP biosynthetic process | 3.25E-02 | 1 | Methanethiol oxidase | B4FAL8 | -2,5 |
| GO:0006790 | sulfur compound metabolic process | 3.37E-02 | 3 | Homocysteine S-methyltransferase 3 | Q9FUM8 | -1,7 |
| | | | | Methanethiol oxidase | B4FAL8 | -2,5 |
| | | | | Anamorsin homolog | B6TB21 | -1,0 |
| GO:0009094 | L-phenylalanine biosynthetic process | 3.71E-02 | 1 | Methanethiol oxidase | B4FAL8 | -2,5 |
| GO:0006817 | phosphate ion transport | 3.71E-02 | 1 | H(+)/Pi cotransporter | Q49B46 | -2,4 |

| | | | | | | |
|---|---|---|---|---|---|---|
| GO:0035970 | peptidyl-threonine dephosphorylation | 4.62E-02 | 1 | Protein-serine/threonine phosphatase | B4FVS8 | -2,0 |
| GO:0071712 | ER-associated misfolded protein catabolic process | 4.62E-02 | 1 | Ubiquitin fusion degradation 1 | C4J8Y3 | -2,1 |
| | | | **3dpi** | | | |
| GO:0009408 | response to heat | 1.66E-03 | 3 | 17.4 kDa class III heat shock protein | B4F9E8 | -3,3 |
| | | | | 17.4 kDa class I heat shock protein 3 | B6TDB5 | -2,9 |
| | | | | Chaperone protein ClpB4 mitochondrial | A0A1D6H2L6 | -1,3 |
| GO:0042542 | response to hydrogen peroxide | 1.97E-03 | 2 | 17.4 kDa class III heat shock protein | B4F9E8 | -3,3 |
| | | | | 17.4 kDa class I heat shock protein 3 | B6TDB5 | -2,9 |
| GO:0051259 | protein complex oligomerization | 3.73E-03 | 2 | 17.4 kDa class III heat shock protein | B4F9E8 | -3,3 |
| | | | | 17.4 kDa class I heat shock protein 3 | B6TDB5 | -2,9 |
| GO:0042026 | protein refolding | 8.29E-03 | 2 | Heat shock 70 kDa protein 5 | K7VJF3 | -4,2 |
| | | | | Chaperone protein ClpB4 mitochondrial | A0A1D6H2L6 | -1,3 |
| GO:0017012 | protein-phytochromobilin linkage | 8.40E-03 | 1 | Phytochrome | Q6XFQ3 | -1,1 |
| GO:0000492 | box C/D snoRNP assembly | 1.67E-02 | 1 | RuvB-like helicase | B6TN62 | -1,0 |
| GO:0009585 | red, far-red light phototransduction | 2.09E-02 | 1 | Phytochrome | Q6XFQ3 | -1,1 |
| GO:0009584 | detection of visible light | 2.09E-02 | 1 | Phytochrome | Q6XFQ3 | -1,1 |
| GO:0009651 | response to salt stress | 2.13E-02 | 2 | 17.4 kDa class III heat shock protein | B4F9E8 | -3,3 |
| | | | | 17.4 kDa class I heat shock protein 3 | B6TDB5 | -2,9 |
| GO:0016925 | protein sumoylation | 3.32E-02 | 1 | SUMO-activating enzyme subunit 1B-2 | B4FTG1 | -1,2 |
| GO:0051716 | cellular response to stimulus | 3.86E-02 | 6 | Peroxidase | A0A1D6N0K3 | -1,2 |
| | | | | Phytochrome | Q6XFQ3 | -1,1 |
| | | | | Protein kinase domain-containing protein | C4IZW2 | -1,8 |
| | | | | Aspartate carbamoyltransferase | C4J935 | -1,2 |
| | | | | Heat shock 70 kDa protein 5 | K7VJF3 | -4,2 |
| | | | | Uncharacterized protein A0A1D6H2L6 | A0A1D6H2L6 | -4,1 |
| GO:0016573 | histone acetylation | 4.54E-02 | 1 | RuvB-like helicase | B6TN62 | -1,0 |
| GO:0016036 | cellular response to phosphate starvation | 4.94E-02 | 1 | Aspartate carbamoyltransferase | C4J935 | -1,2 |
| GO:0006207 | 'de novo' pyrimidine nucleobase biosynthetic process | 4.94E-02 | 1 | Aspartate carbamoyltransferase | C4J935 | -1,2 |

| Proteins with higher abundance | | | | | | |
|---|---|---|---|---|---|---|
| 24 hpi | | | | | | |
| GO:0098869 | cellular oxidant detoxification | 1.04E-05 | 7 | Peroxidase | A0A804UEZ3 | 1,8 |
| | | | | Alpha-dioxygenase 1 | H9BG22 | 1,9 |
| | | | | 4-coumarate--CoA ligase 1;BZIP domain-containing protein | K7UE06 | 0,8 |
| | | | | Peroxidase | C0HHA6 | 1,5 |
| | | | | Peroxidase | C0P4S1 | 2,9 |
| | | | | Peroxidase | D7NLB3 | 1,7 |
| | | | | Uncharacterized protein A0A804QIC5 | A0A804QIC5 | 1,9 |
| GO:0006979 | response to oxidative stress | 4.83E-05 | 7 | Peroxidase | A0A804UEZ3 | 1,8 |
| | | | | Alpha-dioxygenase 1 | H9BG22 | 1,9 |
| | | | | 4-coumarate--CoA ligase 1;BZIP domain-containing protein | K7UE06 | 0,8 |
| | | | | Peroxidase | C0HHA6 | 1,5 |
| | | | | Peroxidase | C0P4S1 | 2,9 |
| | | | | Peroxidase | D7NLB3 | 1,7 |
| | | | | Uncharacterized protein A0A804QIC5 | A0A804QIC5 | 1,9 |
| GO:0042744 | hydrogen peroxide catabolic process | 2.24E-04 | 5 | Peroxidase | A0A804UEZ3 | 1,8 |
| | | | | Peroxidase | C0HHA6 | 1,5 |
| | | | | Peroxidase | C0P4S1 | 2,9 |
| | | | | Peroxidase | D7NLB3 | 1,7 |
| | | | | Uncharacterized protein A0A804QIC5 | A0A804QIC5 | 1,9 |
| GO:0019854 | L-ascorbic acid catabolic process | 6.73E-03 | 1 | Sulfate adenylyltransferase | C0P6J3 | 0,75 |
| GO:1902609 | (R)-2-hydroxy-alpha-linolenic acid biosynthetic process | 6.73E-03 | 1 | Alpha-dioxygenase 1 | H9BG22 | 1,9 |
| GO:0071732 | cellular response to nitric oxide | 6.73E-03 | 1 | Alpha-dioxygenase 1 | H9BG22 | 1,9 |
| GO:0045227 | capsule polysaccharide biosynthetic process | 6.73E-03 | 1 | Sulfate adenylyltransferase | C0P6J3 | 0,75 |
| GO:0050832 | defense response to fungus | 3.95E-02 | 2 | Alpha-dioxygenase 1 | H9BG22 | 1,9 |
| | | | | Allene oxide synthesis4 | B6TT18 | 1,1 |
| GO:0006468 | protein phosphorylation | 2.74E-02 | 5 | Uncharacterized protein B8A0N7 | B8A0N7 | 1,7 |
| | | | | Proline-rich receptor-like protein kinase | B7ZXA1 | 1,0 |

| | | | | | | |
|---|---|---|---|---|---|---|
| | | | | PERK4 | | |
| | | | | Cysteine-rich receptor-like protein kinase 10 | B6UIQ8 | 2,8 |
| | | | | Calmodulin-binding receptor-like cytoplasmic kinase 3 | A0A1D6KAS2 | 1,2 |
| | | | | Uncharacterized protein A0A804UC00 | A0A804UC00 | 1,5 |
| GO:0031408 | oxylipin biosynthetic process | 7.78E-03 | 2 | Alpha-dioxygenase 1 | H9BG22 | 1,9 |
| | | | | Lipoxygenase | C0P840 | 1,3 |
| GO:0001561 | fatty acid alpha-oxidation | 2.01E-02 | 1 | Alpha-dioxygenase 1 | H9BG22 | 1,9 |
| GO:0000256 | allantoin catabolic process | 3.32E-02 | 1 | Sulfate adenylyltransferase | C0P6J3 | 0,75 |
| GO:0006833 | water transport | 8.56E-03 | 2 | Aquaporin PIP1-2 | Q9XF59 | 0,9 |
| | | | | Aquaporin PIP2-2 | A0A1D6GRK5 | 1,9 |
| GO:0006952 | defense response | 3.60E-02 | 4 | Benzoate O-methyltransferase | D9J101 | 1,1 |
| | | | | Uncharacterized protein B8A0N7 | B8A0N7 | 1,7 |
| | | | | Alpha-dioxygenase 1 | H9BG22 | 1,9 |
| | | | | Allene oxide synthesis4 | B6TT18 | 1,1 |
| GO:0010143 | cutin biosynthetic process | 3.97E-02 | 1 | Glycerol-3-phosphate acyltransferase 8 | B6SRR0 | 2,1 |
| GO:0007021 | tubulin complex assembly | 2.01E-02 | 1 | Tubulin-folding cofactor E | A0A1D6H1K6 | 2,0 |
| GO:0007023 | post-chaperonin tubulin folding pathway | 2.01E-02 | 1 | Tubulin-folding cofactor E | A0A1D6H1K6 | 2,0 |
| GO:0051290 | protein heterotetramerization | 4.62E-02 | 1 | Aquaporin PIP1-2 | Q9XF59 | 0,9 |
| GO:0034440 | lipid oxidation | 3.50E-02 | 2 | Alpha-dioxygenase 1 | H9BG22 | 1,9 |
| | | | | Lipoxygenase | C0P840 | 1,3 |
| GO:0000103 | sulfate assimilation | 3.97E-02 | 1 | Sulfate adenylyltransferase | C0P6J3 | 0,75 |
| GO:0019262 | N-acetylneuraminate catabolic process | 1.34E-02 | 1 | Sulfate adenylyltransferase | C0P6J3 | 0,75 |
| GO:0006097 | glyoxylate cycle | 2.67E-02 | 1 | Sulfate adenylyltransferase | C0P6J3 | 0,75 |
| **3dpi** | | | | | | |
| GO:0006556 | S-adenosylmethionine biosynthetic process | 7.18E-05 | 3 | S-adenosylmethionine synthase | B4FAD1 | 1,9 |
| | | | | S-adenosylmethionine synthase | B4FIE9 | 1,8 |
| | | | | S-adenosylmethionine synthase | K7VC35 | 2,5 |
| GO:0006024 | glycosaminoglycan biosynthetic process | 7.20E-04 | 2 | UDP-glucose 6-dehydrogenase | B7ZYX8 | 1,2 |

| | | | | | | |
|---|---|---|---|---|---|---|
| | | | | UDP-glucose 6-dehydrogenase | B6T9P0 | 0,9 |
| GO:0006065 | UDP-glucuronate biosynthetic process | 2.35E-03 | 2 | UDP-glucose 6-dehydrogenase | B7ZYX8 | 1,2 |
| | | | | UDP-glucose 6-dehydrogenase | B6T9P0 | 0,9 |
| GO:0042128 | nitrate assimilation | 3.49E-03 | 2 | Uncharacterized protein A0A804PPM3 | A0A804PPM3 | 1,7 |
| | | | | Ferredoxin--nitrite reductase, chloroplastic | P17847 | 2,5 |
| GO:0010167 | response to nitrate | 6.38E-03 | 2 | Uncharacterized protein A0A804PPM3 | A0A804PPM3 | 1,7 |
| | | | | Ferredoxin--nitrite reductase, chloroplastic | P17847 | 2,5 |
| GO:0009051 | pentose-phosphate shunt, oxidative branch | 8.13E-03 | 2 | Uncharacterized protein A0A804NLI1 | A0A804NLI1 | 2,1 |
| | | | | Glucose-6-phosphate 1-dehydrogenase | B7ZYX8 | 1,7 |
| GO:0009736 | cytokinin-activated signaling pathway | 8.13E-03 | 2 | 4-hydroxy-7-methoxy-3-oxo-3,4-dihydro-2H-1,4-benzoxazin-2-yl glucoside beta-D-glucosidase 1 | P49235 | 1,5 |
| | | | | Beta-glucosidase 17 | A0A1D6MUQ1 | 1,75 |
| GO:0006730 | one-carbon metabolic process | 1.33E-02 | 3 | S-adenosylmethionine synthase | B4FAD1 | 1,9 |
| | | | | S-adenosylmethionine synthase | B4FIE9 | 1,8 |
| | | | | S-adenosylmethionine synthase | K7VC35 | 2,5 |
| GO:0009313 | oligosaccharide catabolic process | 1.44E-02 | 2 | Alkaline/neutral invertase | B8A2U9 | 0,9 |
| | | | | Uncharacterized protein A0A804NLI1 | A0A804NLI1 | 2,1 |
| GO:0009698 | phenylpropanoid metabolic process | 1.49E-02 | 5 | 4-coumarate--CoA ligase 1 | B4FQP4 | 1,0 |
| | | | | Uncharacterized protein A0A804LUI6 | A0A804LUI6 | 1,4 |
| | | | | Acetyl-coenzyme A synthetase | A0A1D6H9J3 | 1,3 |
| | | | | Caffeoyl-CoA O-methyltransferase 1 | Q9XGD6 | 1,3 |
| | | | | Phenylalanine/tyrosine ammonia-lyase | Q8VXG7 | 1,5 |
| GO:0048205 | COPI coating of Golgi vesicle | 1.57E-02 | 1 | ADP-ribosylation factor GTPase-activating protein AGD1 | B4FK00 | 1,4 |
| GO:0036377 | arbuscular mycorrhizal association | 1.57E-02 | 1 | Putative esterase KAI2 | B4FHV3 | 1,9 |
| GO:0009187 | cyclic nucleotide metabolic process | 1.57E-02 | 1 | Cyclic phosphodiesterase | C0P8F5 | 3,3 |
| GO:0009704 | de-etiolation | 1.57E-02 | 1 | Putative esterase KAI2 | B4FHV3 | 1,9 |
| GO:0080167 | response to karrikin | 1.57E-02 | 1 | Putative esterase KAI2 | B4FHV3 | 1,9 |

| | | | | | | |
|---|---|---|---|---|---|---|
| GO:0006308 | DNA catabolic process | 1.57E-02 | 1 | Aspergillus nuclease S(1) | B6T329 | 2,4 |
| GO:0006060 | sorbitol metabolic process | 1.57E-02 | 1 | Uncharacterized protein A0A804LUI6 | A0A804LUI6 | 2,3 |
| GO:0009062 | fatty acid catabolic process | 1.69E-02 | 3 | 3-hydroxyacyl-CoA dehydrogenase | A0A1D6H4X4 | 1,3 |
| | | | | Acyl-coenzyme A oxidase | B4F8X3 | 1,3 |
| | | | | NAD(P)-binding Rossmann-fold superfamily protein | B6T5Z2 | 1,4 |
| GO:0006422 | aspartyl-tRNA aminoacylation | 3.11E-02 | 1 | Aspartyl-tRNA synthetase | B7ZX71 | 1,3 |
| GO:0019745 | pentacyclic triterpenoid biosynthetic process | 3.11E-02 | 1 | Uncharacterized protein A0A804P4K9 | A0A804P4K9 | 1,5 |
| GO:0048658 | anther wall tapetum development | 3.11E-02 | 1 | Protein transport protein SEC23 | K7UW20 | 1,2 |
| GO:0046705 | CDP biosynthetic process | 3.11E-02 | 1 | UMP-CMP kinase | B6T904 | 1,0 |
| GO:0002191 | cap-dependent translational initiation | 3.11E-02 | 1 | Eukaryotic translation initiation factor 3 subunit D | A0A1D6MCY7 | 1,1 |
| GO:0006225 | UDP biosynthetic process | 3.11E-02 | 1 | UMP-CMP kinase | B6T904 | 1,0 |
| GO:0046898 | response to cycloheximide | 3.11E-02 | 1 | Phenylalanine/tyrosine ammonia-lyase | Q8VXG7 | 1,5 |
| GO:0019427 | acetyl-CoA biosynthetic process from acetate | 3.11E-02 | 1 | Acetyl-coenzyme A synthetase | A0A1D6H9J3 | 1,3 |
| GO:0006874 | intracellular calcium ion homeostasis | 3.15E-02 | 2 | Calcium pump1 | K7VI72 | 1,9 |
| | | | | Uncharacterized protein A0A804QQF5 | A0A804QQF5 | 0,9 |
| GO:0009809 | lignin biosynthetic process | 3.15E-02 | 2 | Uncharacterized protein A0A804LUI6 | A0A804LUI6 | 2,3 |
| | | | | Caffeoyl-CoA O-methyltransferase 1 | Q9XGD6 | 1,3 |
| GO:0007264 | small GTPase mediated signal transduction | 3.84E-02 | 2 | Rac-like GTP-binding protein 5 | B4FNE3 | 2,2 |
| | | | | ARF guanine-nucleotide exchange factor GNOM | A0A1D6K8I6 | 1,0 |
| GO:0032259 | methylation | 4.56E-02 | 5 | O-methyltransferase ZRP4 | K7UX80 | 2,0 |
| | | | | Caffeoyl-CoA O-methyltransferase 1 | B6UF45 | 2,5 |
| | | | | Methyltransferase | B4FB47 | 3,1 |
| | | | | Putative methyltransferase | C0PCR5 | 2,0 |
| | | | | Caffeoyl-CoA O-methyltransferase 1 | Q9XGD6 | 1,3 |
| GO:0045048 | protein insertion into ER membrane | 4.62E-02 | 1 | ATPase 100193266 | B4FEF1 | 1,5 |
| GO:0016598 | protein arginylation | 4.62E-02 | 1 | Phenylalanine/tyrosine ammonia-lyase | Q8VXG7 | 1,5 |

| | | | | | | |
|---|---|---|---|---|---|---|
| GO:0035652 | clathrin-coated vesicle cargo loading | 4.62E-02 | 1 | VHS domain-containing protein | A0A1D6MF07 | 1,0 |
| GO:0009626 | plant-type hypersensitive response | 4.62E-02 | 1 | MACPF domain-containing protein CAD1 | C0PEL1 | 0,9 |
| GO:0015976 | carbon utilization | 4.62E-02 | 1 | Carbonic anhydrase | C0P441 | 2,4 |
| GO:0052746 | inositol phosphorylation | 4.62E-02 | 1 | Inositol-pentakisphosphate 2-kinase | K7U7M2 | 1,0 |

**Supplementary Table S4**. Individual proteins with lower and higher abundance, assessed to each KEGG process and pathway at 24 hpi and 3 dpi.

| Pathway group | Pathway subgroup | Pathway | 24 hpi | | | 3 dpi | | |
|---|---|---|---|---|---|---|---|---|
| | | | Protein name | Log | Nr | Protein name | Log | Nr |
| **Metabolism** | Global and overview maps | Biosynthesis of nucleotide sugars | | | | UDP-glucuronate decarboxylase | 0,95 | ↑ 4 |
| | | | | | | UDP-glucose 6-dehydrogenase | 1,2 | |
| | | | | | | Phosphomannomutase | 0,9 | |
| | | | | | | UDP-glucose 6-dehydrogenase | 0,9 | |
| | | Biosynthesis of cofactors | | | | UDP-glucose 6-dehydrogenase | 1,2 | ↑ 8 |
| | | | | | | Phosphomannomutase | 0,9 | |
| | | | | | | S-adenosylmethionine synthase | 1,9 | |
| | | | | | | S-adenosylmethionine synthase | 1,8 | |
| | | | | | | Pyruvate kinase | 0,8 | |
| | | | | | | UDP-glucose 6-dehydrogenase | 0,9 | |
| | | | | | | S-adenosylmethionine synthase | 2,5 | |
| | | | | | | UMP-CMP kinase | 1,4 | |
| | Carbohydrate metabolism | Pentose and glucuronate interconversions | | | | Uncharacterized protein A0A804LUI6 | 2,3 | ↑ 4 |
| | | | | | | UDP-glucose 6-dehydrogenase | 1,2 | |
| | | | | | | UDP-glucose 6-dehydrogenase | 0,9 | |
| | | | | | | Uncharacterized protein A0A804Q287 | 1,8 | |
| | Energy metabolism | Nitrogen metabolism | | | | Carbonic anhydrase | 2,4 | ↑ 4 |
| | | | | | | Uncharacterized protein A0A804PPM3 | 3,1 | |
| | | | | | | Ferredoxin--nitrite reductase chloroplastic | 2,5 | |
| | | | | | | Glutamine synthetase root isozyme 1 | 2,1 | |
| | | alpha-Linolenic acid metabolism | Benzoate O-methyltransferase | 1,1 | ↑ 4 | | | |
| | | | Alpha-dioxygenase 1 | 1,9 | | | | |
| | | | Allene oxide synthesis4 | 1,1 | | | | |
| | | | Lipoxygenase | 1,3 | | | | |
| | | Steroid biosynthesis | | | | Methyltransferase | 3,1 | ↑ 2 |
| | | | | | | Uncharacterized protein A0A804P4K9 | 1,5 | |
| | Amino acid metabolism | Cysteine and methionine metabolism | | | | S-adenosylmethionine synthase | 1,9 | ↑ 5 |
| | | | | | | S-adenosylmethionine synthase | 1,8 | |
| | | | | | | Nicotianamine synthase9 | 4,4 | |
| | | | | | | Tyrosine aminotransferase | 5,1 | |
| | | | | | | S-adenosylmethionine synthase | 2,5 | |

| | | | | | | | | |
|---|---|---|---|---|---|---|---|---|
| | Glycan biosynthesis and metabolism | O-Antigen nucleotide sugar biosynthesis | | | | UDP-glucose 6-dehydrogenas | 1,2 | ↑ 2 |
| | | | | | | UDP-glucose 6-dehydrogenase | 0,9 | |
| | Biosynthesis of other secondary metabolites | Phenylpropanoid biosynthesis | Beta-glucosidase 17 | 1,7 | ↑ 6 | 4-hydroxy-7-methoxy-3-oxo-3,4-dihydro-2H-1,4-benzoxazin-2-yl glucoside beta-D-glucosidase 1, chloroplastic | 1,45 | ↑ 8 |
| | | | Peroxidase | 1,8 | | Beta-glucosidase 17 | 1,75 | |
| | | | Cytochrome P450 84A1 | 0,9 | | Caffeoyl-CoA O-methyltransferase 1 | 1,3 | |
| | | | Peroxidase | 1,7 | | Phenylalanine/tyrosine ammonia-lyase | 1,5 | |
| | | | 4-coumarate--CoA ligase 1;BZIP domain-containing protein | 0,8 | | 4-coumarate--CoA ligase 1 | 1,0 | |
| | | | | | | Shikimate O-hydroxycinnamoyltransferase | 1,4 | |
| | | | Uncharacterized protein A0A804QIC5 | 1,9 | | Peroxidase | 1,7 | |
| | | | | | | Peroxidase | 3,8 | |
| | | Stilbenoid, diarylheptanoid and gingerol biosynthesis | | | | O-methyltransferase ZRP4 | 2,0 | ↑ 3 |
| | | | | | | Caffeoyl-CoA O-methyltransferase 1 | 1,3 | |
| | | | | | | Shikimate O-hydroxycinnamoyltransferase | 1,4 | |
| | | Flavonoid biosynthesis | | | | Caffeoyl-CoA O-methyltransferase 1 | 1,3 | ↑ 3 |
| | | | | | | Chalcone--flavanone isomerase | 2,2 | |
| | | | | | | Shikimate O-hydroxycinnamoyltransferase | 1,4 | |
| | | Neomycin, kanamycin and gentamicin biosynthesis | Uncharacterized protein A0A804PXF7 | 1,9 | ↑ 1 | | | |
| | | Prodigiosin biosynthesis | | | | Uncharacterized protein A0A804NWX6 | -4,1 | ↓ 1 |
| | | Biosynthesis of various secondary metabolites | | | | S-adenosylmethionine synthase | 1,9 | ↑ 5 |
| | | | | | | S-adenosylmethionine synthase | 1,8 | |
| | | | | | | Nicotianamine synthase9 | 4,4 | |
| | | | | | | Tyrosine aminotransferase | 5,1 | |
| | | | | | | S-adenosylmethionine synthase | 2,5 | |
| | Folding, sorting and degradation | Protein processing in endoplasmic reticulum | | | | 17.4 kDa class III heat shock protein | -3,3 | ↓ 3 |
| | | | | | | 17.4 kDa class I heat shock protein 3 | -2,9 | |
| | | | | | | Heat shock 70 kDa protein 5 | -4,2 | |
| **Environmental Information Processing** | Signal transduction | Wnt signaling pathway | | | | Rac-like GTP-binding protein 5 | 2,2 | ↑ 3 |
| | | | | | | Casein kinase II subunit alpha | 0,75 | |
| | | | | | | GLABRA2 expression modulator | 1,5 | |
| | | MAPK signaling pathway | Proline-rich receptor-like protein kinase PERK4 | 1,0 | ↑ 2 | | | |

| | | | | | | | | |
|---|---|---|---|---|---|---|---|---|
| | | | Calmodulin-binding receptor-like cytoplasmic kinase 3 | 1,2 | | | | |
| | | MAPK signaling pathway plant | | | | Pathogenesis-related protein 1 | -2,9 | ↓ 2 |
| | | | | | | Protein kinase domain-containing protein | -1,8 | |
| | | NF-kappa B signaling pathway | Proline-rich receptor-like protein kinase PERK4 | 1,0 | ↑ 2 | | | |
| | | | Calmodulin-binding receptor-like cytoplasmic kinase 3 | 1,2 | | | | |
| | | Plant hormone signal transduction | | | | Pathogenesis-related protein 1 | -2,9 | ↓ 2 |
| | | | | | | Protein kinase domain-containing protein | -1,8 | |
| | | Peroxisome | Superoxide dismutase [Cu-Zn] 2 | -3,9 | ↓ 2 | | | |
| | | | Superoxide dismutase [Cu-Zn] 4AP | -0,8 | | | | |
| | Cellular community | Adherens junction | | | | Rac-like GTP-binding protein 5 | 2,2 | ↑ 2 |
| | | | | | | Casein kinase II subunit alpha | 0,75 | |
| **Organismal systems** | Immune system | Toll-like receptor signaling pathway | Proline-rich receptor-like protein kinase PERK4 | 1,0 | ↑ 2 | | | |
| | | | Calmodulin-binding receptor-like cytoplasmic kinase 3 | 1,2 | | | | |
| | | Toll and Imd signaling pathway | Proline-rich receptor-like protein kinase PERK4 | 1,0 | ↑ 2 | | | |
| | | | Calmodulin-binding receptor-like cytoplasmic kinase 3 | 1,2 | | | | |
| | | Leukocyte transendothelial migration | | | | Rac-like GTP-binding protein 5 | 2,2 | ↑ 1 |
| | Nervous system | Retrograde endocannabinoid signaling | NADH dehydrogenase subunit 2 | -2,7 | ↓ 2 | | | |
| | | | NADH-ubiquinone oxidoreductase chain 1 | -1,9 | | | | |
| | | Neurotrophin signaling pathway | Proline-rich receptor-like protein kinase PERK4 | 1,0 | ↑ 2 | | | |
| | | | Calmodulin-binding receptor-like cytoplasmic kinase 3 | 1,2 | | | | |
| | Development and regeneration | Axon guidance | | | | Rac-like GTP-binding protein 5 | 2,2 | ↑ 2 |
| | | | | | | Actin-depolymerizing factor 10 | 2,4 | |
| | Aging | Longevity regulating pathway – multiple species | Superoxide dismutase [Cu-Zn] 2 | -3,9 | ↓ 2 | Heat shock 70 kDa protein 5 | -4,2 | ↓ 2 |
| | | | NADH-ubiquinone oxidoreductase | -1,9 | | Chaperone protein ClpB4 mitochondrial | -1,3 | |

| | | | |
|---|---|---|---|
| | | | chain 1 |
